# TORC1 modulation in adipose tissue is required for organismal adaptation to hypoxia in *Drosophila*

**DOI:** 10.1101/350520

**Authors:** Byoungchun Lee, Elizabeth C. Barretto, Savraj S. Grewal

## Abstract

Animals often develop in conditions where environmental conditions such as food, oxygen and temperature fluctuate. The ability to adapt their metabolism to these fluctuations is important to ensure normal development and viability. In most animals, low oxygen (hypoxia) is deleterious, however some animals can alter their physiology to thrive under hypoxia. Here we show that TORC1 modulation in adipose tissue is required for organismal adaptation to hypoxia in *Drosophila*. We find that hypoxia rapidly suppresses TORC1 kinase signalling in Drosophila larvae via TSC-mediated inhibition of Rheb. We show that this hypoxia-mediated inhibition of TORC1 specifically in the larval fat body is essential for viable development to adulthood. Moreover, we find that these effects of TORC1 inhibition on hypoxia tolerance are mediated through remodeling of fat body lipid droplets and lipid storage. These studies identify the larval adipose tissue as a key hypoxia sensing tissue that coordinates whole-body development and survival to changes in environmental oxygen by modulating TORC1 and lipid storage.

## INTRODUCTION

Animals often have to grow and survive in conditions where their environment fluctuates. For example, changes in nutrition, temperature or oxygen availability, or exposure to toxins and stress can all impact development. Animals must therefore adapt their physiology and metabolism in response to these environmental challenges in order to ensure proper growth and survival ^1,2^.

In most animals decreases in oxygen are particularly deleterious. Low oxygen (hypoxia) can lead to rapid tissue damage and lethality, and oxygen deprivation is a hallmark of diseases such as stroke and ischemia ^3^. However, some animals have evolved to live in oxygen-deprived conditions and consequently exhibit marked tolerance to hypoxia. For example, birds and aquatic mammals can tolerate extensive periods of low oxygen without incurring any tissue damage ^4,5^. Indeed, some animals show quite remarkable levels of tolerance to oxygen deprivation: brine shrimp embryos have been reported to recover from four years of continuous anoxia ^6^, while the naked mole rat can survive up to 18 minutes of complete oxygen deprivation, a condition that kills laboratory rodents within about one minute ^7^. Understanding how these animals adapt their metabolism to low oxygen may shed light on how to protect tissues from hypoxic damage in disease states.

Drosophila provide an excellent laboratory model system to examine how fluctuations in environmental conditions influence animal development. In particular, there has been extensive work on how nutrient availability influences Drosophila larval development, the main growth period of the life cycle ^8-10^. In nutrient-rich conditions larvae increase in mass approximately 200-fold over four days before undergoing metamorphosis to the pupal stage^11,12^. In contrast, when dietary nutrients are limiting, larvae alter their physiology and metabolism to slow growth and development, and to promote survival. One main regulator of these nutrient-regulated processes in Drosophila is the conserved TOR kinase signalling pathway^13^. TOR exists in two signalling complexes, TORC1 and TORC2, with TORC1 being the main growth regulatory TOR complex ^14^. A conserved signalling network couples nutrient availability to the activation TORC1 to control anabolic process important for cell growth and proliferation^14^. Moreover, studies in Drosophila have been instrumental in revealing non-autonomous effects of TORC1 signalling on body growth. For example, nutrient activation of TORC1 in specific larval tissue such as the fat body, muscle and prothoracic gland, can influence whole animal development through the control of endocrine signalling via insulin-like peptides and the steroid hormone, ecdysone^9,10,15^. In addition, TORC1 regulation of autophagy in the larval fat body is important for organismal homeostasis and survival during periods of nutrient deprivation^16,17^.

Drosophila larvae are also hypoxia tolerant^18-20^. In their natural ecology, Drosophila larvae grow on rotting food rich in microorganisms, which probably contribute to a low oxygen local environment. Even in the laboratory, local oxygen levels are low at the food surface of vials containing developing larvae ^19^. Drosophila have therefore evolved metabolic and physiological mechanisms to respond to and thrive in hypoxic conditions. However, compared to our understanding of the nutrient regulation of growth and homeostasis, considerably less is known about how Drosophila adapt to low oxygen during development. A handful of studies have shown that larval survival in oxygen requires regulation of gene expression by the transcription factors HIF-1 alpha and ERR alpha, and the repressor, Hairy ^21-24^. Developmental hypoxia sensing and signalling has also been shown to be mediated through a nitric oxide/cGMP/PKG signalling pathway^25,26^.

Here we report a role for modulation of the TOR kinase signalling pathway as a regulator of hypoxia tolerance during Drosophila development. In particular, we find that suppression of TORC1 specifically in the larval fat body is required for animals to reset their growth and developmental rate in hypoxia, and to allow viable development to the adult stage. We further show that these effects of TORC1 inhibition require remodelling of lipid droplet and lipid storage. Our findings implicate the larval fat body as a key hypoxia-sensing tissue that coordinates whole animal development and survival in response to changing oxygen levels.

## RESULTS

### Exposure of larvae to hypoxia slows growth and delays development

We began by examining the effect of exposing larvae to hypoxia on their growth and development. We used 5% oxygen as our hypoxia conditions for all experiments in this paper. We allowed embryos to develop in normoxia and then, upon hatching, larvae were either maintained on food in normoxia or transferred to food vials in hypoxia chambers that were perfused with a constant supply of 5%oxygen/95% nitrogen. We found that hypoxia led to reduced larval growth rate and larvae took approximately an extra two days to develop to the pupal stage (Fig 1a). We also found that the hypoxia-exposed animals had a reduced wandering third instar larval weight (Fig 1b) and reduced final pupal size (Fig 1c). We found that exposure of larvae to hypoxia did not alter their feeding behaviour (Suppl Fig 1), suggesting that the decreased growth rate was not simply due to a general reduction in nutrient intake. Together, these data indicate that Drosophila larvae adapt to low oxygen levels by reducing their growth and slowing their development. These data are consistent with previous reports showing that moderate levels of hypoxia (10% oxygen) can also affect final body size ^20^.

**Figure 1.**
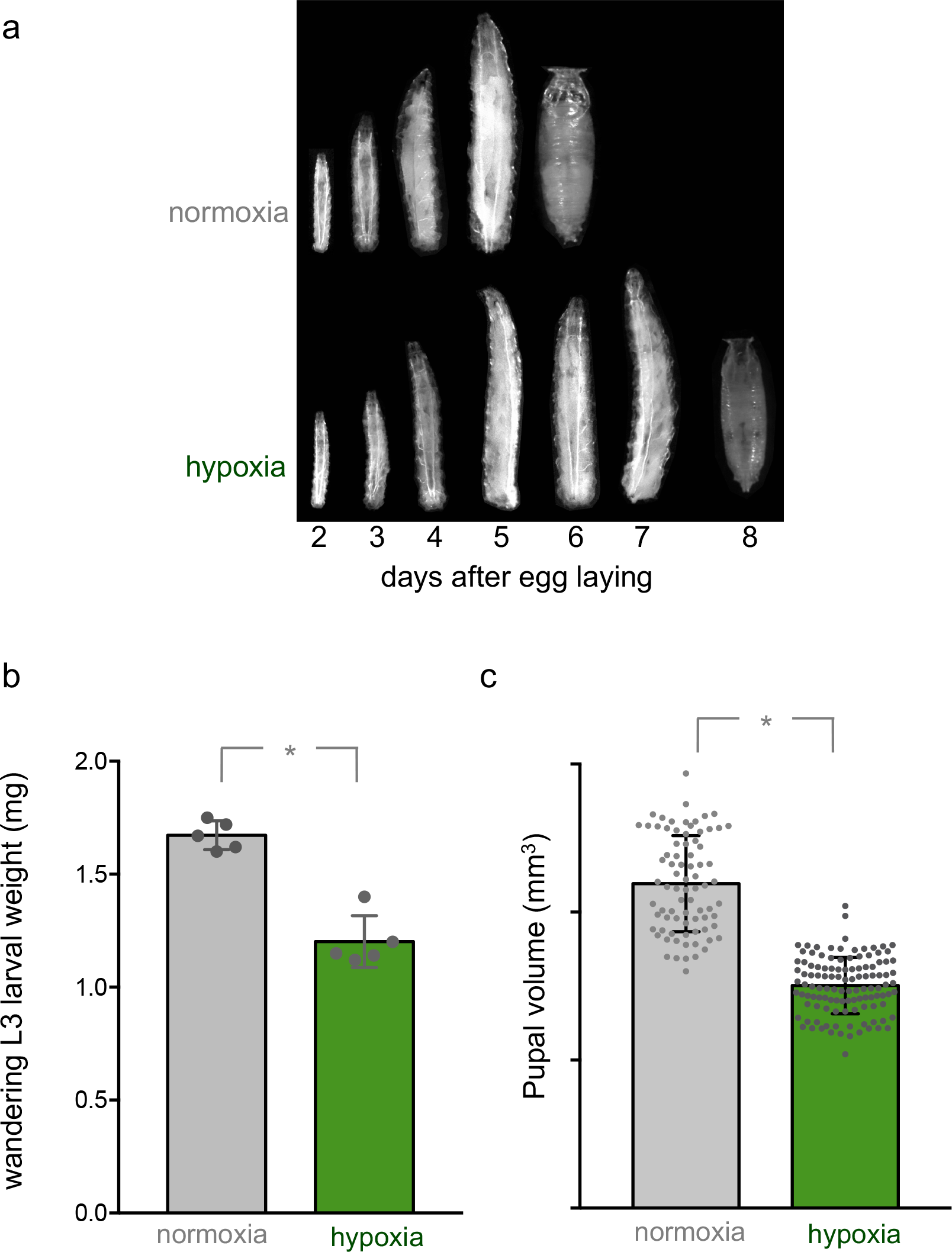
Hypoxia inhibits larval growth and development. **a)** Larvae were hatched in normoxia and then either maintained in normoxia (top images) or transferred to hypoxia (5% oxygen, bottom images). Larvae and pupae were then subsequently imaged on each day following egg hatching. Hypoxia leads to a delay in larval growth and development. **b)** Larvae were hatched in normoxia and then maintained in either normoxia or hypoxia (5% oxygen) until the wandering third instar stage. Larval weights were then measured. Hypoxia lead to a reduction in larval mass. Data are expressed as mean +/− SEM. * = p<0.05. **c)** Larvae were hatched in normoxia and then maintained in either normoxia or hypoxia (5% oxygen) until pupation. Pupal size was then measured. Hypoxia lead to a reduction final pupal size. Data are expressed as mean +/− SEM, * = p<0.05.

### Hypoxia suppresses TORC1 signalling via TSC1/2

The conserved TORC1 kinase signalling pathway is one of the main regulators of tissue and body growth in Drosophila. TORC1 can be activated by dietary nutrients and growth factors such as insulin. Mammalian cell culture experiments have also shown that hypoxia can suppress TORC1 activity^27-30^. We therefore examined whether changes in TORC1 signalling play a role in adaption to hypoxia in Drosophila larvae. We transferred third instar larvae from normoxia to hypoxia and then measured TORC1 activity by western blotting using an antibody that recognizes the phosphorylated form of S6 kinase (pS6K), a direct TORC1 kinase target. We found that hypoxia led to a rapid suppression of whole body TORC1 activity that was apparent within 10-20 minutes of hypoxia exposure (Fig 2a). This suppression persisted when larvae were maintained in hypoxia for longer periods (48 hours, Suppl Fig 2a). We also examined how different levels of oxygen affected TORC1 activity. Third instar larvae were transferred from normoxia to different levels of hypoxia (from 20-1% oxygen) for one hour and then TORC1 activity measured by western blotting for phosphorylated S6K. We found that suppression of TORC1 occurred at 5 and 3% oxygen but remained unchanged at higher (20 and 10%) or lower (1%) levels (Fig 2b). We examined this further by exposing larvae to several different concentrations of oxygen between 1 and 10%, and found that the range within which TORC1 was inhibited was between 2 and 6 % oxygen (suppl Fig 2C). These data indicate that larvae rapidly respond to hypoxia by suppressing TORC1 signalling, and that this response occurs within a specific range of low environmental oxygen rather than simply being triggered below a threshold level of low oxygen.

**Figure 2.**
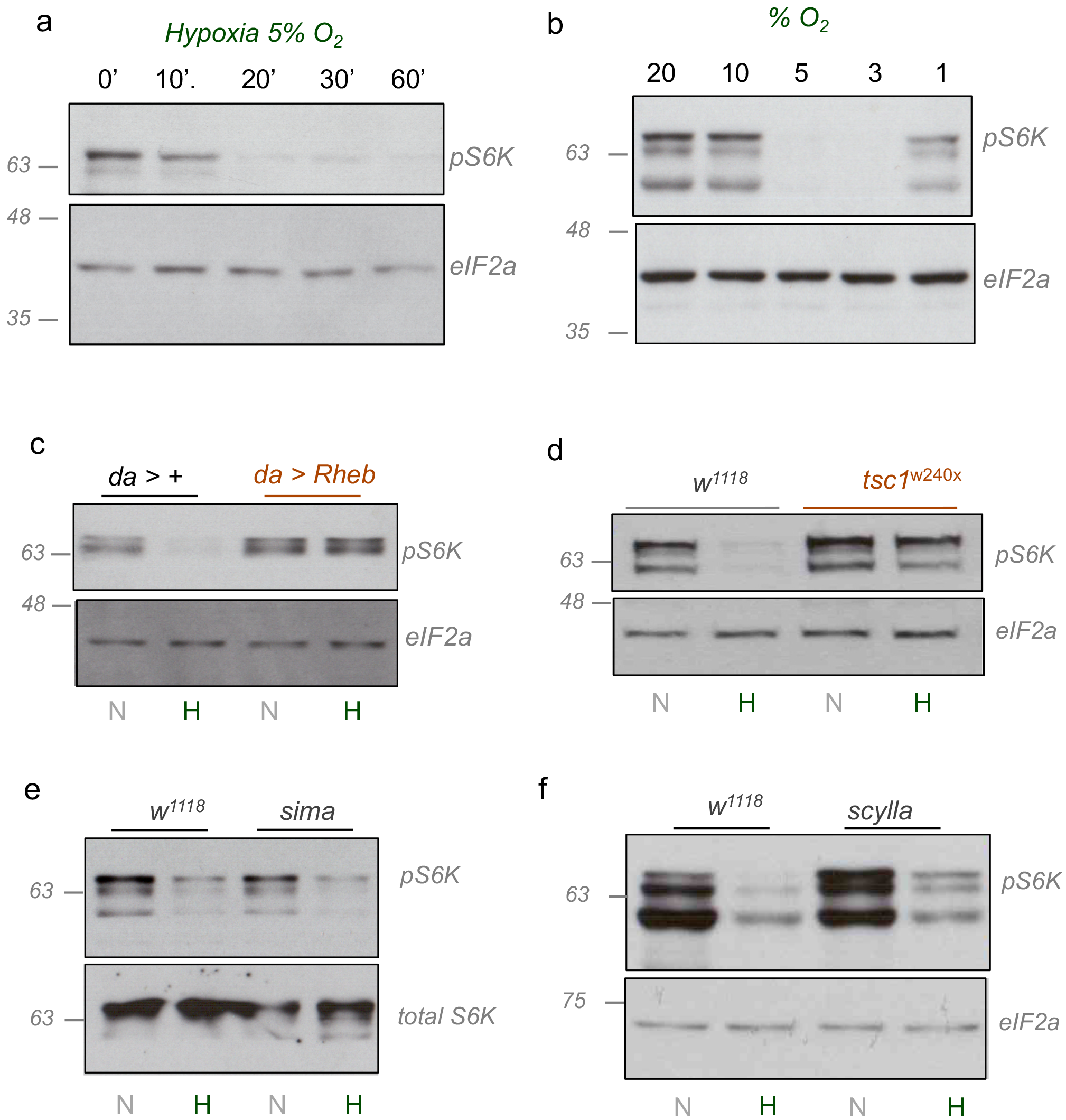
Hypoxia suppresses TORC1 signalling via TSC1/2. **a)** Early third instar larvae were transferred from normoxia to hypoxia (5% oxygen). At the indicated times, larvae were then collected, lysed and processed for SDS-PAGE and western blotting using antibodies to phospho-S6K (pS6K) or total eIF2alpha (eIF2α). Hypoxia led to a rapid suppression of TORC1 signalling. **b)** Early third instar larvae were transferred from normoxia to different levels of hypoxia (20-1% oxygen) for 2hrs. Larvae were then collected, lysed and processed for SDS-PAGE and western blotting using antibodies to phospho-S6K (pS6K) or total eIF2alpha (eIF2α). Suppression of TORC1 signalling was only seen at oxygen concentrations between 1 and 5%. **c)** Control (*da > +)* or Rheb overexpressing (*da>Rheb)* early third instar larvae were either maintained in normoxia (N) or transferred from normoxia to hypoxia (5% oxygen, H) for 2hrs. Larvae were then collected, lysed and processed for SDS-PAGE and western blotting using antibodies to phospho-S6K (pS6K) or total eIF2alpha (eIF2α). Overexpression of Rheb was sufficient to maintain TORC1 signalling in hypoxia larvae at levels seen in normoxic animals. **d)** Control (*w*^*1118*^) or tsc1 mutant (*tsc1*^*Q87X*^) larvae were either maintained in normoxia (N) or transferred from normoxia to hypoxia (5% oxygen, H) for 2hrs. Larvae were then collected, lysed and processed for SDS-PAGE and western blotting using antibodies to phospho-S6K (pS6K) or total eIF2alpha (eIF2α). Hypoxia-mediated suppression of TORC1 signalling required *tsc1* function. **e)** control (*w*^*1118*^) or sima mutant (*sima)* larvae were either maintained in normoxia (N) or transferred from normoxia to hypoxia (5% oxygen, H) for 2hrs. Larvae were then collected, lysed and processed for SDS-PAGE and western blotting using antibodies to phospho-S6K (pS6K) or total S6K. Hypoxia-mediated suppression of TORC1 signalling occurred independently of sima function. **f)** control (*w*^*1118*^) or *scylla* mutant (*scylla*) larvae were either maintained in normoxia (N) or transferred from normoxia to hypoxia (5% oxygen, H) for 2hrs. Larvae were then collected, lysed and processed for SDS-PAGE and western blotting using antibodies to phospho-S6K (pS6K) or total eIF2alpha (eIF2α). Hypoxia-mediated suppression of TORC1 signalling occurred independently of scylla function

We next examined how hypoxia suppresses TORC1 activity. One of the main ways by which TORC1 is activated is through a TSC1/2-Rheb signalling pathway^14^. Rheb is a small G-protein that binds to and activates TOR kinase at lysosomes. TSC2 is a GTPase activating protein, and when bound to its partner TSC1, it inhibits Rheb by converting it from its active GTP-bound state to an inactive GDP-bound state. Several diverse stimuli including nutrients, growth factors and hypoxia have been shown to regulate TSC1/2 function and to control TORC1 activity in mammalian cell culture^14^. We therefore explored a role for TSC1/2 and Rheb in the suppression of TORC1 kinase signalling during larval hypoxia. We found that ubiquitous overexpression of a *UAS-Rheb* transgene (using daughterless-gal4, *da-gal4*) was sufficient to prevent the hypoxia-mediated suppression of TORC1 signalling in larvae (Fig 2c). We also found that tsc1 null mutant (*tsc1*^*Q87X*^) larvae also were unable to suppress TORC1 signalling when exposed to hypoxia (Fig 2d). Together these data indicate that hypoxic exposure in larvae inhibits TORC1 by TSC1/2-mediated suppression of Rheb.

Studies in mammalian cells have described how hypoxia can induce TSC-mediated TORC inhibition via the classic HIF-1 alpha transcription factor. In this mechanism, HIF-1 alpha leads to upregulation of REDD1, an activator of TSC1/2 ^31^. In Drosophila, the homolog of REDD1, Scylla, and its partner protein, Charybdis, have been shown to inhibit TOR and suppress growth ^32^. We therefore examined a role for Sima (the Drosophila HIF-1 alpha homolog) and Scylla/Charybdis in larval hypoxia. However, we found that *sima* mutants still showed a suppression of TORC1 signalling when exposed to hypoxia (Fig 2e).

Similarly, both *scylla* and *charybdis* mutants also showed a similar suppression of TORC1 signalling as control larvae in hypoxia (Fig2f, Suppl Fig 2c). We also explored a potential role for AMPK in hypoxia-mediated TOR regulation. AMPK is activated under hypoxia in mammalian cell culture and can suppress TORC1 signalling, in part by phosphorylating and inhibiting TSC2 ^28,33,34^. However, when we suppressed AMPK by expression of a Gal4-dependent AMPK inverted repeat transgene (*UAS-AMPK IR*), we still saw that hypoxia exposure lead to an inhibition of TORC1 (Suppl Fig 2d). Together, our data suggest that the rapid suppression of TORC1 signalling upon hypoxia exposure in larvae requires TSC1/2 function but is independent of both HIF-1 alpha mediated transcription and AMPK activation.

### Suppression of TORC1 signalling in the fat body is required for adaptation to hypoxia

We next examined whether the suppression of overall TORC1 activity we observed was important for animal adaptation and tolerance to hypoxia during Drosophila development. Our approach was to genetically maintain TORC1 signalling in larvae exposed to hypoxia and then to examine the effects of this manipulation on animal growth, development and survival. To do this, we used the ubiquitous expression of *UAS-Rheb* with *da-Gal4* since we found this condition led to larvae maintaining TORC1 activity under hypoxia (Fig 2c). We compared development in control (*da>+*) vs. Rheb overexpressing (*da>Rheb*) animals that were grown throughout their larval period from hatching in either normoxia or hypoxia. We first found that larval Rheb overexpression had no effect on overall survival to the pupal stage in either normoxia or hypoxia (Fig 3a). We next examined developmental rate by measuring the time to pupation. In normoxic conditions, we found that Rheb overexpression (*da>Rheb*) lead to a slight increase in developmental rate compared to control animals (*da>+*, Fig 3b). When raised in hypoxia, the *da>+* animals had an approximately two-day delay to pupation, and this developmental delay was even further exacerbated in *da>Rheb* animals. (Fig 3b). We also measured effects on overall body size at the end of larval development. We found that *da>Rheb* animals exhibited an increase in both wandering third instar larval weight (Fig 3c) and pupal volume (Fig 3d). These results are consistent with increased growth caused by modest elevation of TORC1 signalling. However, we found that when raised in hypoxia, the increase in size in *da>Rheb* animals was abolished (Fig 3c, **d)**. Given that the *da>Rheb* pupae required an additional ~2 days of larval development to reach the same size as *da >+*, this indicates that the Rheb overexpressing animals actually had a reduced growth rate in hypoxia.

**Figure 3.**
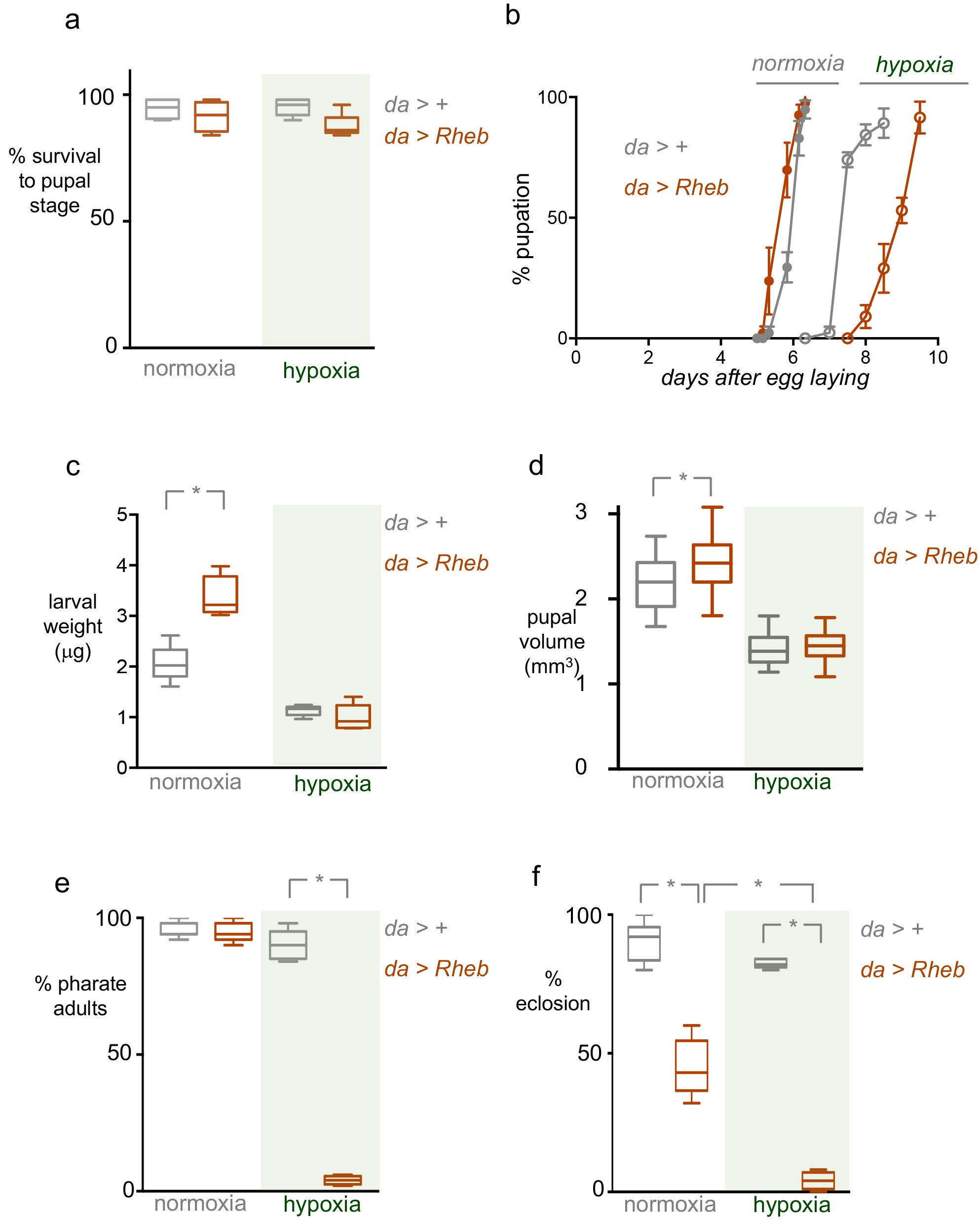
Suppression of TORC1 is required for adaptation to hypoxia. (a-d) Control (*da > +*) or Rheb overexpressing (*da > Rheb*) animals were maintained in either normoxia or hypoxia (5% oxygen) throughout the larval period. **a)** Survival to the pupal stage was measured by calculating the percentage or larvae that developed to pupae for each experimental condition. **b)** The rate of larval development was measured by calculating the percentage of animals that progressed to the pupal stage over time. Maintaining TORC1 signalling in larvae during hypoxia led to a further delay in pupation. **c)** Weights of wandering third instar larvae were measured for each experimental condition. Maintaining TORC1 signalling in hypoxia did not increase larval growth. Data are presented as box plots (25%, median and 75% values) with error bars indicating the min and max values. N=6 groups of larvae per condition. **d)** Pupal volume was calculated for each experimental condition. Maintaining TORC1 signalling in larvae during hypoxia did not increase final pupal size. Data are presented as box plots (25%, median and 75% values) with error bars indicating the min and max values. N >100 pupae per condition. * = p<0.05. e, **f)** Control (*da > +*) or Rheb overexpressing (*da>Rheb*) animals were maintained in either normoxia or hypoxia (5% oxygen) throughout the larval period, before being returned to normoxia at the pupal stage. The % of animals that developed to **e)** pharate adults, or **f)** eclosed adults were then calculated. Maintaining TORC1 signalling in larvae during hypoxia lead to a subsequent lethality during pupal development. Data are presented as box plots (25%, median and 75% values) with error bars indicating the min and max values. N= 6-8 independent groups of animals (50 animals per group) per experimental condition. * = p<0.05.

Finally, we examined how maintaining TORC1 activity during larval development in hypoxia affects subsequent survival to adulthood. For these experiments, we maintained animals in either normoxia or hypoxia throughout their larval period and then switched them to normoxia and monitored their development. We first saw that animals carrying either the *da>Gal4* (*da>+)* or UAS-Rheb (*+>Rheb*) transgenes alone had no effect on viability in either normoxia or hypoxia (Suppl Fig 3a). We found that both *da>+* and *da>Rheb* animals grown in normoxia as larvae showed normal development to the pharate adult stage. Similarly, *da>+* animals grown in hypoxia as larvae also showed no significant change in development to pharate adults. In contrast, *da>Rheb* animals that were maintained in hypoxia during their larval period showed a marked lethality at the pupal stage with few animals developing to pharate adults (Fig 3d). When we further examined adult eclosion, we again saw that *da>Rheb* animals that were maintained in larval hypoxia showed almost complete lethality, but in this case the *da>Rheb* animals raised in normoxia also showed a reduction in eclosion, albeit to a much lesser extent than their hypoxia-raised counterparts. We repeated our Rheb overexpression experiments with a second independent *UAS-Rheb* transgene and we observed similar, but slightly weaker effects, where *da>Rheb* animals grown in hypoxia as larvae showed a significant decrease in survival to adult stage compared to *da>+* animals (Suppl Fig 3b).

Taken together, these experiments using ubiquitous expression of Rheb to maintain TORC1 signaling indicate that suppression of TORC1 is required for larvae to reset their development and growth rate in hypoxic environments, and for subsequent viable development to the adult stage.

The adaptation to hypoxia may reflect a cell-autonomous requirement for each cell to sense low oxygen and inhibit TORC1 to promote overall development and survival. Alternatively, hypoxia may modulate TORC1 in one particular tissue to control overall body growth and development. A precedent for this is the nutrient regulation of larval physiology and growth. For example, nutrient-dependent changes in TORC1 signalling in specific tissues such as the fat body or prothoracic gland can control whole animal growth and development through non-autonomous effects on endocrine signaling. In this manner, one tissue functions as a sensor of environmental stimuli to coordinate whole body responses. To examine a potentially similar role in hypoxia sensing, we examined whether TORC1 suppression in a specific tissue was required for hypoxia tolerance in developing Drosophila. To do this we again took the approach of expressing a *UAS-Rheb* transgene to maintain TORC1 signaling under hypoxia, but this time we restricted Rheb expression to specific larval tissues. We chose to examine effects on hypoxia tolerance by maintaining animals in either normoxia or hypoxia during their larval period and then measuring survival to eclosion. We tested Gal4 drivers that express in the fat body (*r4-Gal4*), neurons (*elav-Gal4*), the intestine (*MyoIA-Gal4*), the prothoracic gland (*P0206-Gal4*) and the muscle (*dmef2-Gal4*). We found the most dramatic effects were seen with fat-specific expression of Rheb: *r4>Rheb* animals grown in hypoxia during their larval stage showed a significant decrease in adult survival compared to *r4>+* control animals (Fig 4a). However, in contrast to ubiquitous expression of Rheb, we found that fat body restricted expression did not delay larval development in hypoxia - *r4>Rheb* animals developed slightly faster to the pupal stage in both normoxia and hypoxia compared to control (*r4>+*) animals. Also, *r4>Rheb* animals showed no significant change in final pupal size compared to *r4>+*) animals. When we performed similar experiments with expression of Rheb in either neurons, intestine or prothoracic gland we saw no effect on viability (Fig 4b-d). Animals expressing Rheb in muscle (*dmef2>Rheb*) did show reduced adult survival when grown in hypoxia as larvae, however they also showed reduced survival in normoxia, making the effects on hypoxia tolerance difficult to interpret (Fig 4e).

**Figure 4.**
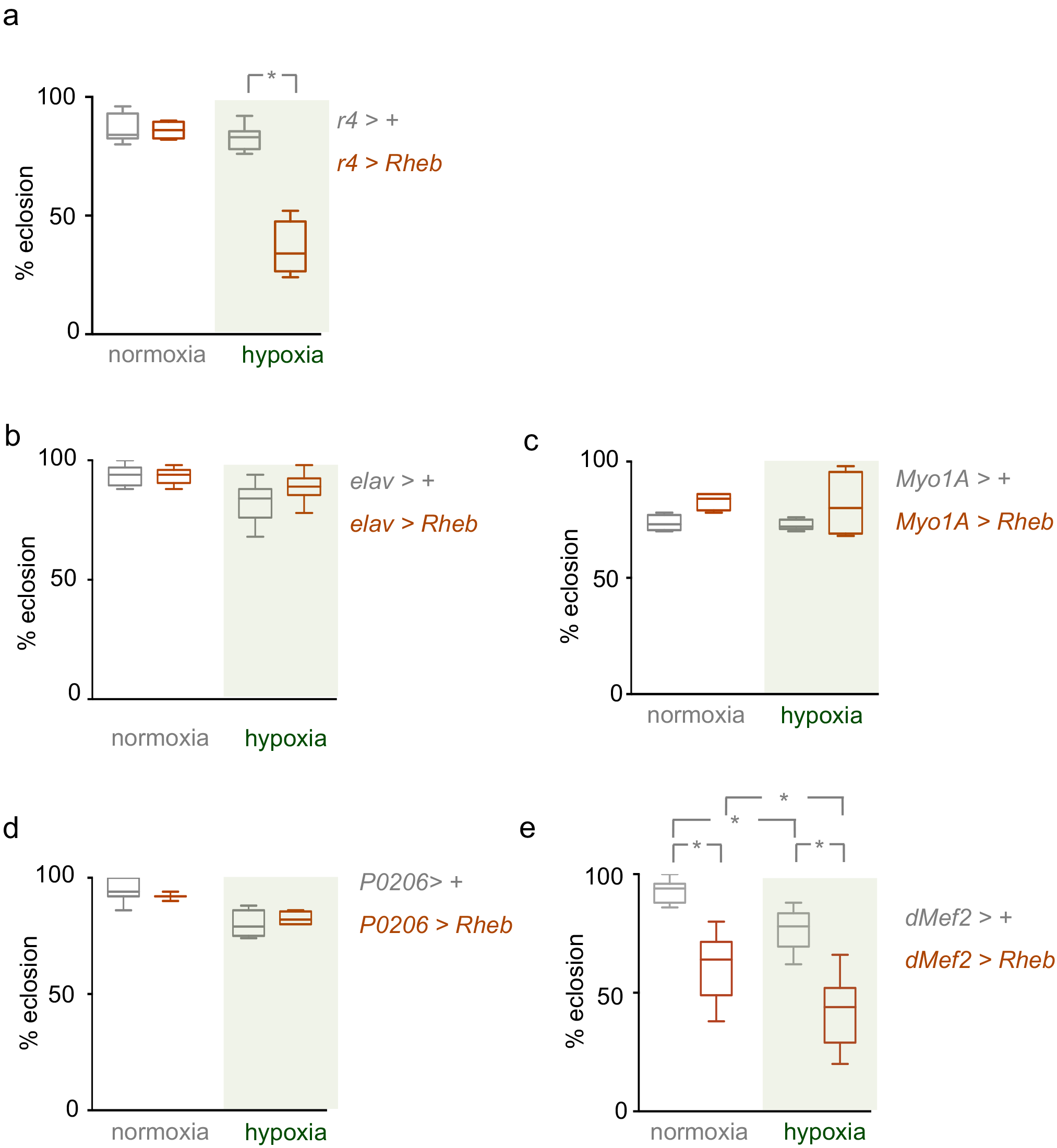
Suppression of TORC1 in the fat body is required for adaptation to hypoxia. **a)** Control (*r4 > +*) larvae or larvae overexpressing Rheb in the fat body (*r4 >Rheb*) were maintained in either normoxia or hypoxia (5% oxygen) throughout the larval period and then were returned to normoxia at the beginning of the pupal stage. The percentage of animals that eclosed as viable adults was then measured. Animals expressing Rheb in the fat body and exposed to hypoxia as larvae showed a significant decrease in adult survival. Data are presented as box plots (25%, median and 75% values) with error bars indicating the min and max values. N= 4-7 groups of animals (50 animals per group) per experimental condition. * = p<0.05. **b-e)** Rheb was overexpressed in larval neurons (**c**, *elav* > *Rheb*), the intestine (**d**, *MyolA > Rheb*), the prothoracic gland (**e**, *P0206 > Rheb*) or muscle (**f**, *dmef2* > *Rheb*). Animals were hatched in normoxia and then maintained throughout the larval period in either normoxic or hypoxic conditions, before being returned to normoxia at the pupal stage. The percentage of animals that developed to the adult stage was calculated for each experimental condition. Control animals carried the Gal4 transgene alone. Data are presented as box plots (25%, median and 75% values) with error bars indicating the min and max values. N= 4-8 independent groups of animals (50 animals per group) per experimental condition. * = p<0.05.

These results suggest that the larval fat body is an important hypoxia sensing tissue that responds to low oxygen by suppressing TORC1 activity to ensure subsequent viable development. We therefore focused our attention on understanding how reduced TORC1 signaling in the fat body contributes to hypoxia tolerance.

### Suppression of TORC1 signalling in the fat body leads to increases in lipid droplet size and lipid storage

We next examined how reduction of TORC1 signaling in the larval fat body contributes to normal organismal development and survival in hypoxia. The role of the fat body as a coordinator of overall body physiology and development has been best studied in the context of altered dietary nutrients. In particular, when larvae are starved of nutrients the fat body mobilizes stored sugars and lipids in order to maintain circulating levels of these nutrients and support tissue homeostasis ^9, 10^. Upon starvation, fat body cells also rapidly engage autophagy to promote organismal survival ^17^. We therefore examined whether these changes are associated with exposure to low oxygen. We first examined autophagy since this is a well-studied conserved process known to be induced by TORC1 inhibition. We subjected early third instar larvae to hypoxia for six hours and then stained fat bodies with LysoTracker Red to visualize lysosomes and late stage autophagosomes as an indicator of autophagy. We also stained fat bodies from larvae maintained in normoxia and from larvae subjected to six hours of nutrient starvation, a condition known to induce autophagy. We found that fat bodies from normoxic animals showed little staining with LysoTracker Red, while starved fat bodies showed a marked increase in LysoTracker Red punctae, consistent with induction of autophagy (Fig 5). In contrast, we saw little or no LysoTracker Red punctae in fat bodies from larvae exposed to hypoxia for six hours (Fig 5). Even longer hypoxia exposure (24 hours) also did not induced autophagy.

**Figure 5.**
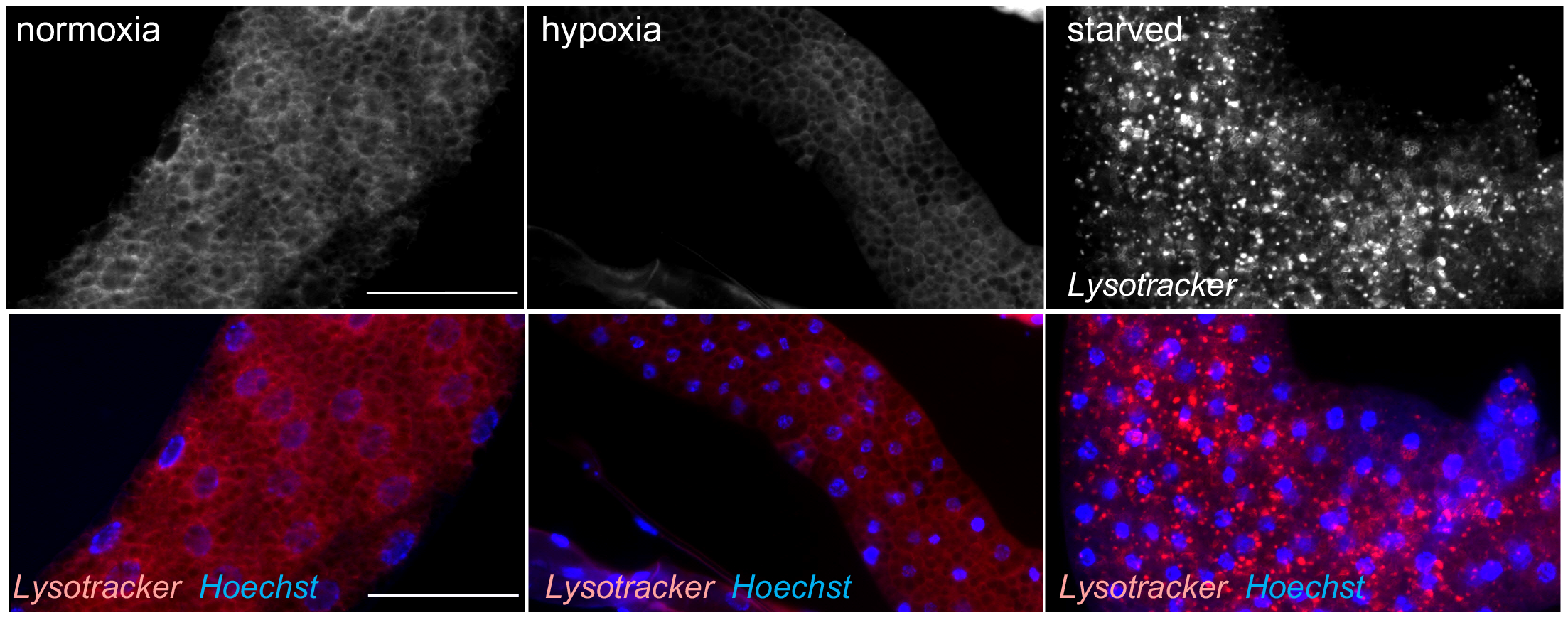
Hypoxia does not induce autophagy. Early third instar larvae were maintained in normoxia (left panels), transferred to hypoxia for 6 hours (middle panels), or starved on PBS for 6 hours (right panels). Fat bodies were then dissected, stained with LysoTracker and then imaged. Red = LysoTracker; blue = Hoechst DNA stain. Scale bar = 50μm. Starvation, but not hypoxia, induced autophagy in the fat body.

We then explored effects on lipid metabolism. In the fat body, triacylglycerol (TAGs) are stored within large lipid droplets. These lipid stores then can be mobilized under starvation conditions to supply a source of free fatty acid for beta-oxidation and other metabolic processes required for homeostasis ^35^. We observed that when larvae were raised in hypoxia they showed a noticeable change in fat body morphology, which became less opaque in appearance as has been reported previously ^36^. When we examined the fat bodies under light microscopy we saw an increase in cytoplasmic lipid droplet size (Fig 6a). We examined this phenotype in more detail by using Nile Red to stain the neutral lipids that compose these cytoplasmic lipid droplets. When we transferred second instar larvae to hypoxia for two days we observed a significant increase lipid droplet diameter compared to larvae maintained in normoxia for the same period (Fig 6b, c). This effect on lipid droplets was opposite to that seen in larvae that were starved of all nutrients for two days (PBS only), which exhibited a marked decrease in lipid droplet size (Fig 6b). Instead, the hypoxia phenotype was similar to animals that were transferred to a sugar-only diet for two days. These results indicate that the effects of hypoxia on lipid droplet size are opposite to those seen in nutrient-deprivation and suggest that under hypoxia larvae may increase TAG levels through increase synthesis from dietary sugars. To measure TAG levels more quantitatively, we raised larvae from hatching in either normoxia or hypoxia and then measured whole-body TAG levels using a colorimetric assay. We found that hypoxic animals exhibited approximately a two-fold increase on total TAG levels when corrected for total larval weight (Fig 6d). We additionally used a previously described sucrose solution buoyancy assay to estimate larval lipid content ^37,38^. In this assay groups of isolated wandering third instar larvae are mixed with increasing concentrations of a sucrose solution and the percentage of larvae floating at each concentration is measured. Using this approach, we found that hypoxic larvae were more buoyant than larvae growth in normoxia, consistent with an increase in lipids as a proportion of total body mass (Fig 6e). Altogether, these results indicate that hypoxia induces a remodelling of lipid droplet and an increase in total lipid storage.

**Figure 6.**
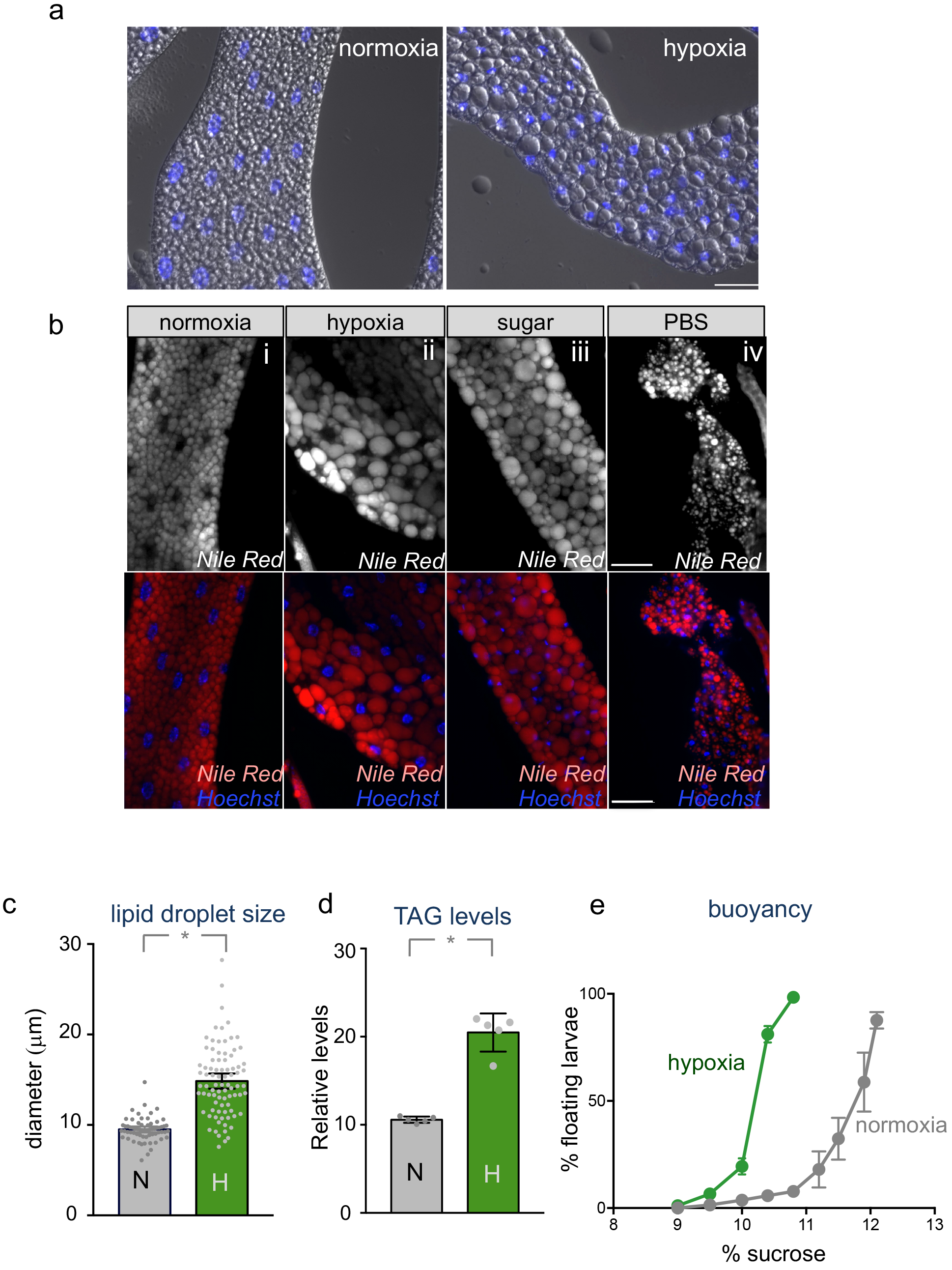
Hypoxia alters lipid levels and lipid storage. **a)** Larvae were hatched in normoxia and then maintained throughout the larval period in either normoxic or hypoxic conditions. Fat bodies from third instar larvae were then imaged using DIC microscopy. Blue = Hoechst DNA stain. Hypoxia lead to an increase in lipid droplet size in fat body cells. Scale bar = 50μm **b)** w1118 larvae were grown in normoxia for until 72 hrs after egg laying at which point they were transferred to one of four experimental conditions for 48 hours: (i) normoxia, (ii) hypoxia, (iii) sugar only diet, (iv) complete starvation (PBS) diet). Fat bodies were then dissected, stained with Nile Red and then imaged. Red + Mile Red, blue = Hoechst DNA dye. Hypoxia lead to an increase in lipid droplet size, similar to a sugar only diet. Scale bar = 50μm **c)**. Lipid droplet size from fat bodies represented in Bi and Bii were measured and presented as mean diameter +/-SEM. **d)** Larvae were hatched in normoxia and then maintained throughout the larval period in either normoxic or hypoxic conditions. Total TAG levels from third instar larvae from both experimental conditions were then measured. Hypoxia exposure lead to an increase in larval total TAG levels. Data are presented as mean +/−SEM. **e)** Larvae were hatched in normoxia and then maintained throughout the larval period in either normoxic or hypoxic conditions. Larval lipid content was then estimated in wandering third instar larvae using an assay, which measures the percentage of larvae that float in increasing amounts of a sucrose solution. Hypoxia-exposed larvae floated at lower concentrations of sucrose compared to normoxic larvae, indicative of a higher lipid content in hypoxic larvae.

We next examined whether these changes in lipid metabolism occurred as a consequence of reduced TORC1 activity. To test this, we generated GFP-marked fat body *tsc1* mutant cell clones. As we previously described, loss of TSC1 completely reversed the hypoxia-mediated suppression of TORC1 signaling. Hence, we examined these *tsc1* mutant fat body cells to see if they still showed the hypoxia-mediated changes in lipid droplets. We induced clones during mitosis in the embryo and then when the animals hatched we transferred them to hypoxia for their entire larval development. When we dissected and examined the fat bodies from third instar larvae using DIC microscopy, we observed the hypoxia increase in lipid droplet size in all non-GFP cells (Fig 7). However, the *tsc1* mutant cells showed no increase in lipid droplet size. Instead they maintained the small lipid droplet morphology typical of normoxic animals at the same stage even though the animals had been grown in hypoxia for several days (Fig 7). These data indicate suppression of TORC1 signalling is required of the hypoxia-mediate remodelling of lipid storage.

**Figure 7.**
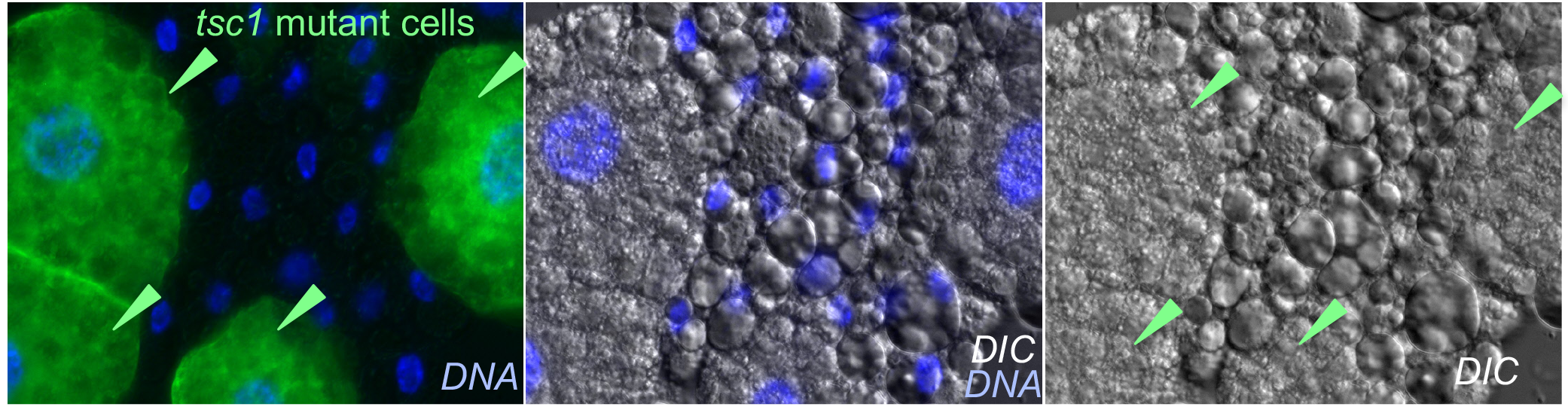
TORC1 suppression is required for hypoxia-induced modulation of lipid storage. The MARCM system was used to generate GFP-marked *tsc1* mutant cell clones in the fat body. Hatched larvae were then either maintained in normoxia or transferred to hypoxia. At the third instar stage, larval fat bodies were fixed, dissected and mounted on coverslips. Fat bodies were then imaged using DIC microscopy to visualize lipid droplets. Blue = Hoechst DNA dye.

### Reorganization of lipid metabolism is required for hypoxia tolerance

We next examined whether the changes in lipid storage caused by the hypoxia-mediated suppression of fat body TORC1 signaling was important for development and survival. To do this we used genetic knockdown of Lsd2, a Drosophila perilipin homolog^39-41^. Lsd2 is a protein associated with the surface of lipid droplets that is necessary for normal lipid droplet formation. We used expression of an inverted repeat (IR) to Lsd2 (*UAS-lsd2 IR*) to specifically knockdown Lsd2 in the fat body using the *r4-Gal4* driver. When we did this and then transferred animals to hypoxia for two days, we found that the large lipid droplet phenotype seen in control (*r4>+*) animals was blocked when Lsd2 levels were reduced (*r4>lsd2 IR*; Fig 8a). We then explored how this inhibition of lipid droplet size affected tolerance to hypoxia. We maintained *r4>+* and *r4>lsd2 IR* larvae in hypoxia from larval hatching to pupation, and then switched them back to normoxia and monitored viability to adult stage. We found that the *r4>lsd2 IR* showed a significant reduction in survival compared to *r4>+* control animals (Fig 8b). To confirm this effect, we also examined a previously reported *lsd2* mutant allele (*lsd2*^*KG00149*^). These *lsd2* mutants are viable and show normal development when grown on normal laboratory food in normoxia. However, when we maintained these *lsd2* mutants in hypoxia throughout their larval period, they showed a marked reduction in survival to adult stage compared to control (*w*^1118^) animals (Fig 8c). These results indicate that the increase in lipid droplet size caused by reduced TORC1 is required for organismal adaptation to hypoxia.

**Figure 8.**
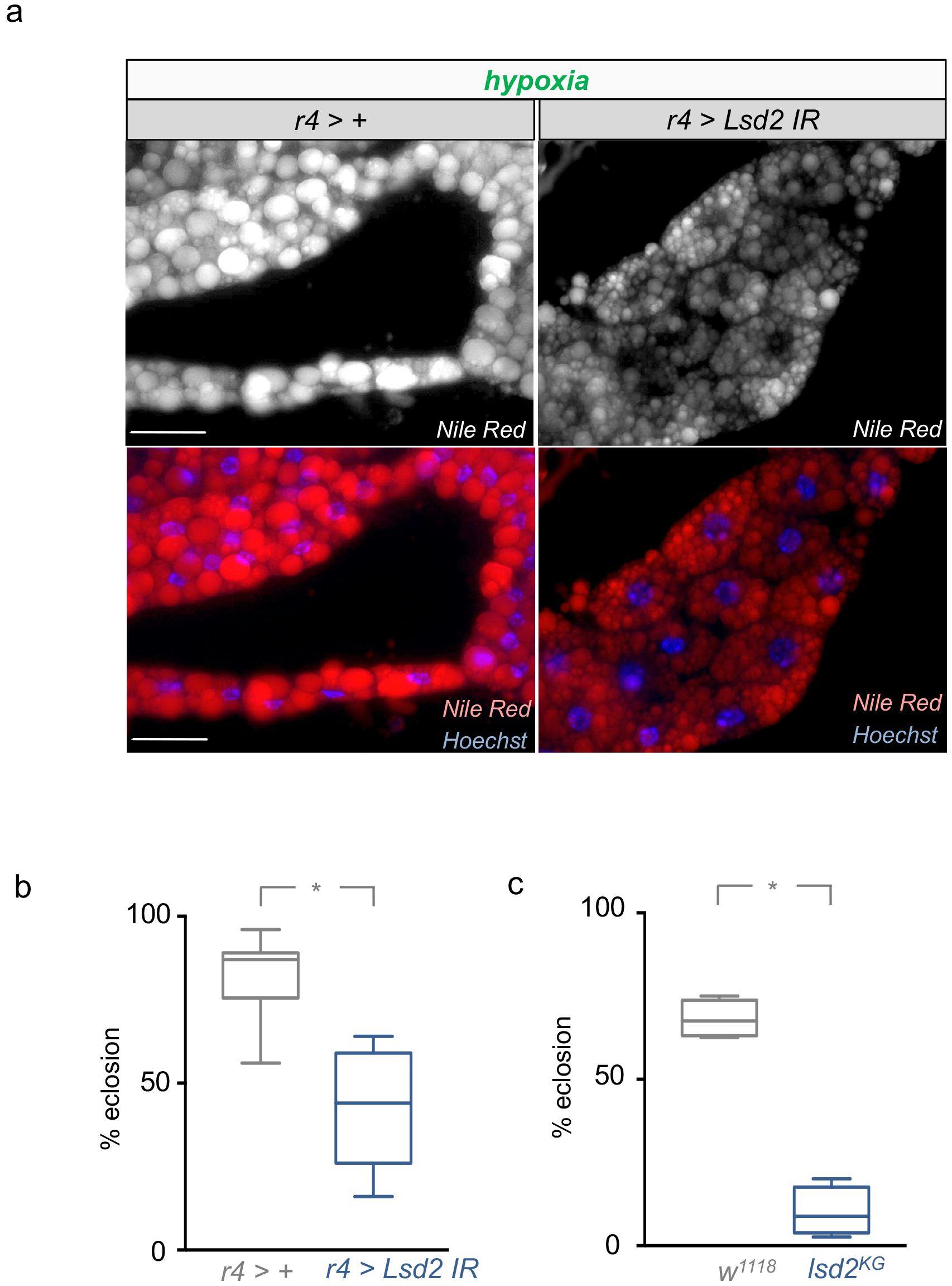
Reorganization of lipid droplets is required for adaptation to hypoxia. **a)** Control larvae (*r4 > +*) or larvae expressing an inverted repat RNAi transgene to eLsd2 (*r4 > Lsd2 IR*) were transferred to hypoxia at 72hrs. After 48 hours of hypoxia, fat bodies were dissected and stained with Nile Red. RNAi-mediated knockdown of Lsd2 inhibited the increase in lipid droplet size caused by hypoxia. Scale bar = 50μm **b)** Control larvae (*r4 > +*) or larvae expressing an inverted repeat RNAi transgene to Lsd2 (*r4 > Lsd2 IR*) were maintained in hypoxia throughout the larval period from hatching to pupation. Animals were then returned to normoxia and the percentage of eclosed adults counted. RNAi-mediated knockdown of Lsd2 lead to decreased adult survival. Data are presented as box plots (25%, median and 75% values) with error bars indicating the min and max values. N= 4-8 groups of animals (50 animals per group) per experimental condition. * = p<0.05. **c)** Control (*w*^*1118*^) or *lsd2*mutant (*lsd2KG*) larvae were maintained in hypoxia throughout the larval period from hatching to pupation. Animals were then returned to normoxia and the percentage of eclosed adults counted. *Lsd2* mutant animals show decreased survival to adult stage. Data are presented as box plots (25%, median and 75% values) with error bars indicating the min and max values. N= 4-8 independent groups of animals (50 animals per group) per experimental condition. * = p<0.05.

## DISCUSSION

In this paper, we explored how Drosophila are able to tolerate hypoxia. A central finding of our work is that when larvae are exposed to low oxygen, the fat body serves as a key hypoxia sensor that mediates changes in physiology to ensure viable organismal development. This hypoxia sensor role is mediated through inhibition of TORC1 signaling and reorganization of lipid storage. This function of the fat body as a hypoxia sensor is reminiscent of the role of the fat body is coordinating whole body physiology responses to changes in dietary nutrients ^9,10,17,42,43^. As we find in hypoxia, these nutrient effects are also dependent on modulation of TORC1 activity and they can exert both metabolic and endocrine effects to control growth and development. These studies and our findings in hypoxia, emphasize how the fat body functions a sentinel tissue to detect changes in environmental conditions and to buffer the internal milieu from these changes. Moreover, while most work on hypoxia has focused on studying cells in culture^27,44^, our findings emphasize the importance of non-cell autonomous mechanisms in controlling how animals adapt to low oxygen.

Inhibition of TORC1 in larvae exposed to hypoxia occurred rapidly and, interestingly, only in response to a specific range of low oxygen (˜2-6%). At <2% oxygen and lower, the response to hypoxia is very different compared to exposure to 5% oxygen that was used in this study - larvae crawl away from the food and eventually undergo complete movement arrest, which can be reversed within minutes of return to normoxia. Larvae can only tolerate this level of low oxygen (<2%) for a few hours before dying. Since this low oxygen hypoxic response is different to the behaviour of larvae at 5% oxygen (which maintain their feeding and growth) it may also rely on qualitatively different changes in hypoxia sensing and signaling that do not involve suppression of TORC1. The hypoxia-mediated inhibition of TORC1 that we found required TSC1/2 but was independent of two main mechanisms defined in mammalian cell culture experiments - induction of REDD1 by the well-studied HIF-1 alpha transcription factor or by activation of AMPK. Although the Drosophila homolog of REDD1, Scylla, was previously shown to be sufficient to inhibit TORC1 ^32^, we found that it was not necessary. Indeed, analysis of the REDD1 mutant mouse also showed that in certain tissues, hypoxia-mediated repression of TORC1 was also REDD1-independent ^34^. A previous report in cell culture showed that upon different stresses including hypoxia, TSC2 could translocate to the lysosome and inhibit Rheb activation of TORC1 ^45^. Therefore, upon hypoxia exposure in larvae, the TSC1/2 complex may rapidly re-localize to inhibit TORC1 function. The mechanism that could drive this (or any other potential mechanism of TORC1 inhibition) must be triggered rapidly in response to hypoxia in larvae. Given the importance of oxygen as an electron acceptor in the electron transport chain in the mitochondria, it is plausible that the rapid sensing of low oxygen in larvae occurs as a result of altered mitochondrial activity. Two potential hypoxia effectors in this scenario are induction of reactive oxygen species or alterations in the levels of mitochondrial metabolites. For example, both 2-hydroxyglutarate and alpha ketoglutarate - metabolites in the TCA cycle - can alter TORC1 activity ^46,47^ and both can be induced by hypoxia in mammalian cell culture experiments ^48^. How these changes could then subsequently lead to an increase in TSC1/2 function and/or localization to inhibit TORC1 remains to be determined.

We found that the protective effect of lowering adipose TORC1 signaling on hypoxia involved the reorganization of lipid metabolism. Hypoxic larvae increase their fat body lipid droplet size and their total proportion of whole body TAGs. This response is very different from starvation, a stress that also inhibits TORC1, where lipid droplet size is markedly reduced. This has been reported to be due to upregulation of lipases such as the ATGL lipase, Brummer ^49-51^. This mechanism provides a way to generate a source of free fatty acids for beta-oxidation to maintain homeostasis and to allow survival under scarce nutrient conditions. In our experiments, the hypoxia induced lipid droplet phenotypes were similar to those seen when larvae were switched to a sugar only diet. One interpretation of this result is that under hypoxia larvae mobilize dietary glucose toward new TAG lipid synthesis and storage. Hence, the large lipid droplet and increased TAG levels seen in hypoxia may be primarily due to new TAG synthesis rather than a suppression of lipolysis. Our data also suggest that the effect of TORC1 inhibition on lipid storage differs between the stress stimuli that suppress TORC1 (hypoxia vs starvation). Indeed, we also saw that decreased TORC1 activity in hypoxia did not trigger autophagy in the fat body in the same way that it does upon nutrient deprivation.

One result we found interesting was that the suppression of fat body TORC1 and altered larval lipid storage was not necessary for viable larval development under hypoxia, but was required for subsequent development in the pupal stage to produce viable adults. During the pupal stage, tissues undergo metamorphosis to establish the adult body. Since this is also a non-feeding stage of the life cycle, the energy required to fuel these extensive tissue rearrangements in pupae must therefore come from stored nutrients. It has been calculated that the lipid stores provide 90% of this energy ^52^. Our findings suggest that pupae may be more dependent on these lipid stores after a period of prior larval hypoxia. Hence failure to maintain these stores, either by preventing TORC1 inhibition (Rheb overexpression) or genetic disruption of lipid droplet formation (Lsd2 knockdown), lead to reduced viability in hypoxia, while having no effect on normal development in normoxia. It is also possible that the requirement for altered lipid stores may reflect a role for lipid droplets beyond simply providing a usable energy source ^53^. A pertinent example is a report describing how increases in glial lipid droplets in larvae were important for maintaining neuroblast cell proliferation in larvae exposed to hypoxia or oxidative stress ^54^. In this case, the lipid droplets were required to play an antioxidant role to buffer neurons from ROS-induced damage. Mammalian cancer cells in culture have also been shown to accumulate lipid droplets in low oxygen, an effect that is important to promote their survival and tumorigenic phenotypes in mouse models ^55^. Cancer cells with high levels of TORC1 activity have also been shown be dependent on exogenous fatty acids for their survival in hypoxic conditions ^56,57^. Hence, the lipid droplet phenotypes we observed may be important for ensuring cell and tissue viability in pupal stages independent of any role in energy production.

In conclusion, our studies presented here pinpoint the Drosophila fat body as a key hypoxia sensing tissue that ensures viable animal development in low oxygen. We suggest that, given the importance of the fat body as a regulator of adult *Drosophila* physiology, this hypoxia-sensing role may be a general mechanism of low oxygen tolerance throughout *Drosophila* life.

## METHODS

### *Drosophila* stocks

Flies were raised on standard medium containing 150 g agar, 1600 g cornmeal, 770 g Torula yeast, 675 g sucrose, 2340 g D-glucose, 240 ml acid mixture (propionic acid/phosphoric acid) per 34 L water and maintained at 25°C, unless otherwise indicated. The following fly stocks were used:

*w*^*1118*^*, tsc1*^*Q87X*^*/TM6B* ^*58*^*, sima*^*07607*^*/TM3, Ser, GFP* ^*21*^*, scylla/TM3,Ser,GFP* ^*32*^*, charybdis*^*180*^*/TM3* ^*32*^*, Ser,GFP, lsd2*^*KG00149*^*, UAS-AMPK RNAi (VDRC), UAS-Rheb (Bloomington Stock Centre), UAS-Lsd2 RNAi* (Bloomington Stock Centre, #), *da-Gal4, r4-Gal4, P0206-Gal4, Elav-Gal4, MyoIA-Gal4, dmef2-GAL4*.

For all GAL4/UAS experiments, homozygous GAL4 lines were crossed to the relevant UAS line(s) and the larval or adult progeny were analyzed. Control animals were obtained by crossing the relevant homozygous GAL4 line to flies of the same genetic background as the particular experimental UAS transgene line.

### Hypoxia exposure

For all hypoxia experiments (except for those shown in Figure 2b) Drosophila were exposed to 5% oxygen. This was achieved by placing vials containing Drosophila into an airtight glass chamber into which a mix of 5%oxygen/95% nitrogen continually flowed. Flow rate was controlled using an Aalborg model P gas flow meter. Alternatively, for some experiments Drosophila vials were placed into a Coy Laboratory Products in vitro O_2_ chamber that was maintained at fixed oxygen levels of 1-20% (depending on the nature of the experiment: See Figure 2b and Suppl Fig 2b) by injection of nitrogen gas.

### Preparation of protein extracts

*Drosophila* larvae were lysed with a buffer containing 20 mM Tris-HCl (pH 8.0), 137 mM NaCl, 1 mM EDTA, 25 % glycerol, 1% NP-40 and with following inhibitors 50 mM NaF, 1 mM PMSF, 1 mM DTT, 5 mM sodium ortho vanadate (Na_3_VO_4_) and Protease Inhibitor cocktail (Roche Cat. No. 04693124001) and Phosphatase inhibitor (Roche Cat. No. 04906845001), according to the manufacturer instructions.

### Western blots and antibodies

Protein concentrations were measured using the Bio-Rad Dc Protein Assay kit II (5000112). Protein lysates (15 μg to 30μg) were resolved by SDS-PAGE and electro transferred to a nitrocellulose membrane, subjected to Western blot analysis with specific antibodies, and visualized by chemiluminescence (enhanced ECL solution (Perkin Elmer). Primary antibodies used in this study were: anti-phospho-S6K-Thr398 (Cell Signalling Technology #9209), anti-eIF2 alpha (AbCam #26197), anti-S6K a gift from Aurelio Teleman), anti-actin (Sant Cruz Biotechnology, # sc-8432). Secondary antibodies were purchased from SantaCruz Biotechnology (sc-2030, 2005, 2020). For experiments looking at TORC1 activity, either total eIF2 alpha, actin or total S6K levels were used as loading controls because the levels of these proteins were unaffected by hypoxia.

### Measurement of *Drosophila* development, growth and survival

#### Development timing to pupal stage

newly hatched larvae were collected at 24hr AEL and placed in food vials (50 larvae per vial). The number of pupae was counted each day. For each experimental condition, a minimum of 5 replicates was used to calculate the mean percentage of pupae per time point.

#### Larval growth

newly hatched larvae were collected at 24hr AEL and placed in food vials (50 larvae per vial) and then maintained in either normoxia or hypoxia. Larvae were then imaged on each day of development using a Zeiss Discovery.V8 Stereomicroscope with Axiovision imaging software. For Fig1A, images of larvae were taken on different days. The larval images were then cropped and arranged as shown in the figure.

#### Larval Weight

Third instar larvae were washed in PBS, dried thoroughly on paper and then weighed in groups of ten using a microbalance.

#### Pupal Volume

Pupae were imaged using a Zeiss Discovery.V8 Stereomicroscope with Axiovision imaging software. Pupal length and width were measured and pupal volume was calculated using the formula, volume = 4/3π(L/2)(l/2)^2^

### LysoTracker and Nile red staining

Fat bodies were dissected from larvae and incubated in either Nile Red (1:50,000 dilution of a 10% stock in DMSO, ThermoFisher Scientific, N1142) or LysoTracker (1:1000, Thermofisher Scientific, L7528) for 10 mins on glass slides. Fat bodies were then immediately imaged using a Zeiss Observer Z1 microscope using Axiovision software.

### Lipid measurements

Groups of ten larvae were washed in PBS, dried on filer paper and then weighed. Total TAG levels were determined as described in detail in ^59^. The calculated TAG levels were then corrected for larval weight to give a measure of total TAG levels per microgram of larval weight. The buoyancy assay was carried out as described in detail in ^38^.

### Statistical analyses

Data were analyzed by Students t-test, or two-way ANOVA followed by post-hoc students t-test where appropriate. All statistical analysis and data plots were performed using Prism software. In all figures, statistically significant differences are presented as: * p<0.05.

## ACKNOWLEDGEMENTS

We thank Paula Bellosta, Aurelio Teleman, Hugo Stocker, Ernst Hafen, Bruce Edgar and Iswar Hariharan for the gift of reagents and fly stocks. Stocks obtained from the VDRC, the NIG-Fly Stock Centre, Kyoto, Japan and the Bloomington Drosophila Stock Center (NIH P40OD018537) were used in this study. This work was supported by a CIHR operating grant and a NSERC Discovery grant to S.S.G. E.C.B. was supported by Alberta Innovates Health Solutions Graduate Studentship.

---

**Supplemental Figure 1.**
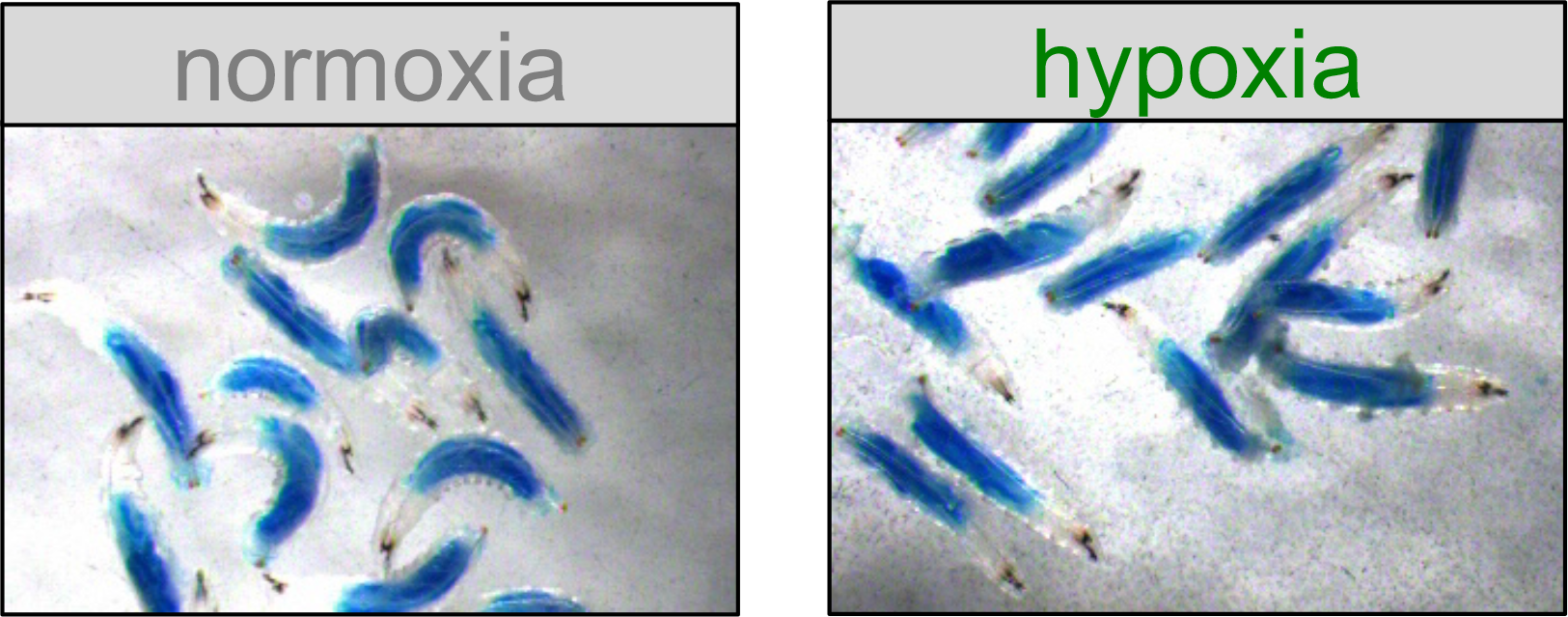
Animals exposed to hypoxia maintain normal food intake. Early third instar larvae were placed into vials containing food mixed with blue food dye. The vials were then either maintained in normoxia or placed into hypoxia for one hour. Larvae were then imaged using a Zeiss Stereomicroscope. The larvae exposed to hypoxia showed similar levels of food intake as larvae maintained in hypoxia.

**Supplemental Figure 2.**
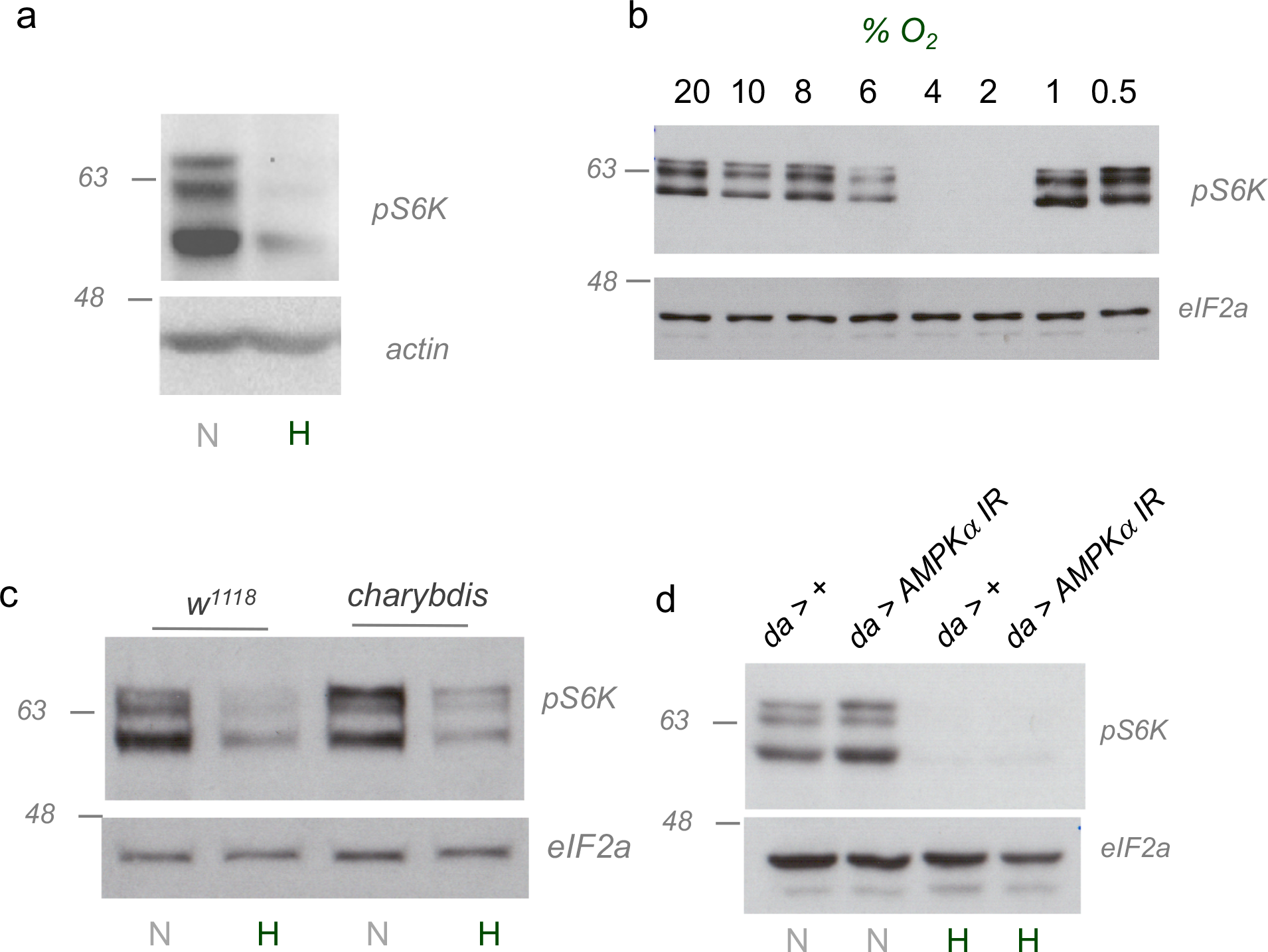
Hypoxia-dependent suppression of TORC1 signaling is independent of AMPK and Charybdis. **a)** Early third instar larvae were either maintained in normoxia or transferred from normoxia to hypoxia (5% oxygen) for 40 hours. Larvae were then collected, lysed and processed for SDS-PaGe and western blotting using antibodies to phospho-S6K (pS6K) or actin. **b)** Early third instar larvae were transferred from normoxia to different levels of hypoxia (20-1% oxygen) for 2hrs. Larvae were then collected, lysed and processed for SDS-PAGE and western blotting using antibodies to phospho-S6K (pS6K) or total eIF2alpha (eIF2α). **c)** control (*w*^*1118*^) or *charybdis* mutant larvae were either maintained in normoxia (N) or transferred from normoxia to hypoxia (5% oxygen, H) for 2hrs. Larvae were then collected, lysed and processed for SDS-PAGE and western blotting using antibodies to phospho-S6K (pS6K) or total eIF2alpha (eIF2α). **d)** Control (*da > +*) or AMPKα RNAi overexpressing (*da >AMPKα IR*) early third instar larvae were either maintained in normoxia (N) or transferred from normoxia to hypoxia (5% oxygen, H) for 2hrs. Larvae were then collected, lysed and processed for SDS-PAGE and western blotting using antibodies to phospho-S6K (pS6K) or total eIF2alpha (eIF2α).

**Supplemental Figure 3.**
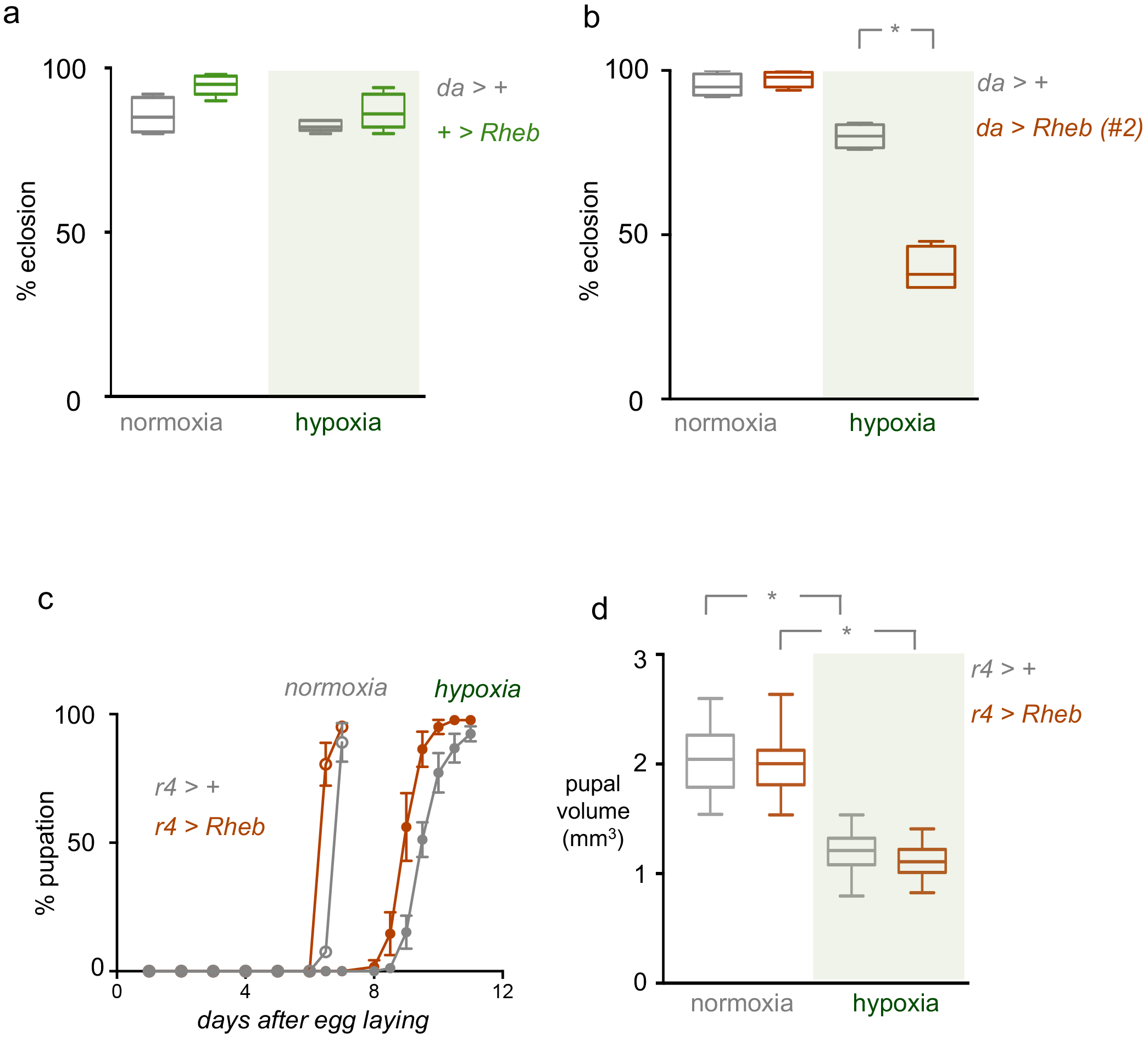
Suppression of TORC1 is required for adaptation to hypoxia. **a)** Larvae carrying either the *da-Gal4 (da>+)* or the *UAS-Rheb (+>Rheb)* transgene were maintained in either normoxia or hypoxia (5% oxygen) throughout the larval period and then were returned to normoxia at the beginning of the pupal stage. The percentage of animals that eclosed as viable adults was then measured. Data are presented as box plots (25%, median and 75% values) with error bars indicating the min and max values. N= 4-7 groups of animals (50 animals per group) per experimental condition. **b)** Control (*da > +*) larvae or larvae overexpressing a second independent UAS-Rheb transgene (*da>Rheb#2*) were maintained in either normoxia or hypoxia (5% oxygen) throughout the larval period and then were returned to normoxia at the beginning of the pupal stage. The percentage of animals that eclosed as viable adults was then measured. Animals expressing Rheb and exposed to hypoxia as larvae showed a significant decrease in adult survival. Data are presented as box plots (25%, median and 75% values) with error bars indicating the min and max values. N= 4-7 independent groups of animals (50 animals per group) per experimental condition. * = p<0.05. **c)** Control (*r4 > +*) larvae or larvae overexpressing Rheb in the fat body (*r4 >Rheb*) were maintained in either normoxia or hypoxia (5% oxygen) throughout the larval period. The rate of larval development was measured by calculating the percentage of animals that progressed to the pupal stage over time. Maintaining TORC1 signalling in the larval fat body did not reverse the hypoxia-mediated delay in larval development. Data points represent mean +/− SEM, N>5 groups of animals per experimental condition. **d)** Control (*r4 > +*) larvae or larvae overexpressing Rheb in the fat body (*r4 >Rheb*) were maintained in either normoxia or hypoxia (5% oxygen) throughout the larval period. Pupal volumes were then measured for each experimental condition. Data are presented as box plots (25%, median and 75% values) with error bars indicating the min and max values. N >100 pupae per condition. * = p<0.05.

